# Reactive and predictive processes during unpredictable driving collisions in virtual reality: an exploratory immersive study with multimodal neurophysiological monitoring

**DOI:** 10.64898/2026.09.24.754154

**Authors:** Cédric Cannard, Demet Yeşilba

## Abstract

Anticipatory neurophysiological activity before unpredictable emotional events has been reported for decades, but the effects are small, difficult to replicate, and obtained almost exclusively in stripped-down laboratory paradigms. Whether they reflect genuine predictive processes or methodological artifacts remains debated; one rarely tested possibility is that such settings mute them. We examined anticipation in an immersive virtual reality (VR) driving simulation whose collision outcomes were assigned by a quantum random number generator, while electroencephalography (EEG), photoplethysmography, electromyography, electrooculography and inertial signals were recorded from a single wireless dry-electrode headset built into the VR display. Sixteen participants passively observed 120 driving sequences ending in a collision or no collision. Length-matched pre- and post-stimulus windows were analyzed identically with mass-univariate hierarchical general linear models under permutation cluster correction, plus machine-learning classification across held-out participants. Two post-stimulus clusters differentiated the conditions, the largest peaking at 451.6 ms (d = −1.35), alongside a broadband power increase robust to baseline modeling; post-stimulus EEG was decoded from held-out participants with up to 93.8% accuracy. No time-domain effect or decodable information preceded stimulus onset, but a broadband spectral difference did, surviving preregistered controls for expectation bias (run length), time-on-task build-up and ocular or facial-muscle artifact: directionally consistent in 13 of the 15 participants with verified trial order, with pre- and post-stimulus power coupled in all 16. Heart rate differentiated the conditions neither alone nor when added to the classifier. The study was preregistered; the data, analysis code and an open-source EEGLAB plugin are publicly available.

## 1. Introduction

The human brain continuously predicts forthcoming sensory events, a core proposal of predictive processing accounts (Clark, 2013; Friston, 2010; Hohwy, 2013). Anticipation is best characterized when cues signal what or when something will happen. The contingent negative variation (CNV), a slow frontocentral negativity, indexes temporal and motivational anticipation in cued paradigms (Walter et al., 1964; Tecce, 1972; Brunia and van Boxtel, 2001) and tracks learned temporal regularities even without motor preparation (Mento et al., 2013). Its amplitude grows when the inter-stimulus interval is stable and is larger before emotionally salient stimuli (Poli et al., 2007; van Boxtel and Böcker, 2004). How the brain responds to, and whether it prepares for, genuinely unpredictable and highly salient events in ecologically valid settings remains unclear.

Most electrophysiological work on threat processing uses laboratory stimuli: static images or tones on a screen, usually in one sensory modality. These paradigms are well controlled, but they lack the temporal and multisensory structure of real hazards. If defensive preparation evolved for events that unfold in time and engage several senses, a brief, repeated laboratory stimulus may elicit only a weak version of the response. The same work typically records a single physiological modality, leaving cortical and autonomic indices of preparation to be inferred from separate studies, or not at all. Virtual reality (VR) can preserve experimental control while restoring ecological validity (Parsons, 2015; Reggente et al., 2018), and collisions in immersive driving simulations are particularly suitable: they have abrupt, multisensory onset and clear survival relevance.

Electroencephalography (EEG) studies in driving contexts have begun to examine these events, with limitations. Li et al. (2022) recorded EEG across four stages of pedestrian collision avoidance and found broad increases in delta, theta, alpha and beta power. Active braking confounded threat processing with movement preparation, band-power analysis lacked temporal resolution, and one event per participant precluded trial-level analysis. Wang et al. (2025) examined directed connectivity while participants watched driving videos and predicted collisions, reporting more stable beta-band connectivity in experienced drivers. Their stimuli were flat-screen videos, only correctly predicted trials were analyzed, and collision timing varied naturally, leaving temporal expectancy uncontrolled.

A separate literature asks whether physiology before randomly assigned, emotionally salient events differs by upcoming event type without predictive cues. A meta-analysis of 26 studies reported a small but consistent effect (weighted d = 0.21, 95% CI [0.15, 0.27]; Mossbridge et al., 2012). Effects of this size require large samples to replicate reliably, and they have been obtained almost exclusively in the stripped-down settings described above: brief, repetitive, emotionally mild stimuli on a screen, with at most one physiological modality recorded. If anticipatory activity is a preparatory response shaped by real-world threat, paradigms so far removed from it may be measuring its weakest version, a parsimonious alternative to attributing the difficulty of replication to chance alone. This was a primary motivation for the present study. This literature also identified an artifact that can mimic such effects: the expectation bias or gambler’s fallacy (Dalkvist et al., 2002). When stimuli are drawn with replacement, participants may implicitly expect an arousing event after a run of neutral ones, and averaging across sequences can create a spurious pre-stimulus difference. Using a true hardware random number generator with equiprobable categories effectively eliminates this bias (Dalkvist et al., 2002; Tressoldi et al., 2015).

Several EEG studies have reported pre-stimulus differentiation between upcoming categories. McCraty et al. (2004) found frontal amplitude differences before emotional and neutral pictures, Radin and Lobach (2007) found occipital differences before light flashes, and Radin et al. (2011) found CNV-like activity in experienced meditators, although their causal elliptic filter entailed substantial phase distortion. Duma et al. (2017) observed a CNV-like negativity about 1000 ms before crash trials in a driving simulation with randomly presented crashes. A second preregistered study from the same group used faces and sounds and did not reach significance in its confirmatory analyses, which the authors attributed to conservative correction across a high-density montage (Duma et al., 2019). A further unpublished study from the group reported larger anticipatory activity over occipital regions before faces than sounds, and over right auditory regions before sounds than faces (Duma et al., unpublished results).

Two methodological issues recur across this literature and motivate the present design. First, zero-phase (bidirectional) filters, applied by default in common software packages, smear post-stimulus activity backwards into the pre-stimulus period and can generate entirely artifactual anticipatory effects (Acunzo et al., 2012; Rousselet, 2012; Widmann and Schröger, 2012). Bilucaglia et al. (2021) illustrate the risk: a zero-phase finite impulse response (FIR) filter of order 16,500 at 500 Hz has an impulse response extending about 16.5 s in each direction, so their decoded 1000 ms window almost certainly contained post-stimulus information shifted backwards. Causal minimum-phase filters cannot do this by construction. Second, baseline correction can contaminate pre-stimulus epochs when the inter-stimulus interval is too short for activity to return to baseline. When conditions are compared with pairwise permutation contrasts, as here, shared drift or offset cancels in the difference, so baseline correction is unnecessary and costs signal-to-noise ratio (Maess et al., 2016; Delorme, 2023).

The present study addresses these issues together. Participants passively observed unpredictable collisions in an immersive VR driving simulation, combining ecological salience with control over event timing and randomization. Trial sequences were generated with a quantum random number generator (qRNG) at equiprobable rates, so the interval preceding a trial carried no information about its outcome, and the passive design avoided motor-preparation confounds. EEG was filtered causally and analyzed without baseline correction, using symmetric pre- and post-stimulus windows.

Our primary aim was to characterize the post-stimulus response to unexpected collisions at the single-trial level, which prior VR-EEG collision work has not done. Secondary aims were to test whether pre-stimulus activity differs between upcoming collision and no-collision trials under these controls, and to compare cortical and cardiac responses to the same events within one session, headset and analysis pipeline, so that the two modalities could be compared directly rather than across studies.

Six hypotheses were preregistered. H1 held that post-stimulus EEG would differ between collision and no-collision events, with an early sensory response to the unexpected onset within approximately 100–150 ms, followed by extended differences over roughly 300–1000 ms as the collision unfolded. H2, the primary anticipatory hypothesis, held that pre-stimulus EEG would differentiate upcoming collision from no-collision events in the second preceding onset, despite the absence of any predictive cue. H3 held that such an effect, if predictive rather than a CNV, would remain stable across early, middle and late blocks rather than growing with learning. H4 held that the magnitude of pre-stimulus differentiation would correlate positively with the magnitude of the post-stimulus response, as time-symmetric accounts require. H5 held that pre-stimulus differentiation would be stronger in participants with greater experience in domains demanding sustained attention and anticipation. H6 held that a classifier trained on pre-stimulus features would discriminate upcoming events above chance. H3 and H4 were evaluated on the time-frequency clusters (Sections 2.7 and 3.4).

## 2. Materials and methods

### 2.1 Preregistration

The study was preregistered at https://osf.io/xuw34 prior to data analysis, as a confirmatory investigation targeting N = 63. Data collection ended at N = 18, and electrodermal activity (EDA) and pupillometry did not yield usable data (Section 2.4). We therefore report all analyses as exploratory, draw no confirmatory inference from any result, positive or null, and treat effect sizes as provisional: the achieved sample was not powered to detect effects of the magnitude reported in the pre-stimulus literature. The registered target corresponded to 80% power for a paired effect of 0.36 (two-tailed α = .05); with the 16 participants analyzed, a single paired comparison at the same α reaches 80% power only for effects of d ≥ 0.75 (d ≥ 0.81 for the 14 participants with usable heart-rate data), before any correction for multiple comparisons. The analyses also deviated from the registered protocol in several respects; every deviation and its rationale is documented in an update to the original registration on the same record, which should be read alongside this report. Quality-control decisions about sensor integrity and data retention were made without reference to experimental condition, none of these decisions was informed by a condition contrast, and all analysis code is available at the repository listed under Data availability.

### 2.2 Participants

Participants were recruited from the general adult population through online advertisements, mailing lists, and word of mouth. Inclusion criteria were age ≥ 18 years, ability to travel to the laboratory, no known screen sensitivity, and ability to participate safely in a VR driving task. Exclusion criteria were uncorrected visual impairment, any physical or psychological condition interfering with task performance, and a history of severe motion sickness or trauma related to car accidents. All participants gave written informed consent and received $30 compensation. All study activities were approved and overseen by the Institutional Review Board (IRB) at the Institute of Noetic Sciences (IORG#0003743).

Data collection ended after 18 participants. Of these, one was excluded for retaining only 27 artifact-free trials per condition and one for insufficient EEG signal quality, yielding a final EEG sample of N = 16 (10 female, 6 male). A separate quality screen was applied to the cardiac data (Section 2.9), giving N = 14 for that analysis; sensor-specific screening was used because signal quality of the EEG and photoplethysmography (PPG) channels was independent within participants. Mean age was 56.8 years (SD = 14.7, range 27–76) and mean formal education 17.1 years (SD = 3.6). Self-rated experience (0–10) was: driving M = 7.0 (SD = 1.3), sports M = 3.6 (SD = 1.8), video games M = 3.6 (SD = 3.0), meditation M = 3.9 (SD = 2.5). Fourteen participants (87.5%) reported believing in intuition and two (12.5%) responded “I don’t know”. Personality was assessed with the Ten-Item Personality Inventory (TIPI; Gosling et al., 2003), administered on a five-point agreement scale rather than the instrument’s standard seven-point scale, so trait scores range from 1 to 5 and are not comparable with published TIPI norms. Items were presented grouped by trait rather than in the instrument’s original interleaved order. It was completed by 14 of the 16 participants (trait means 2.9 to 4.2 on the 1–5 scale).

### 2.3 Design and virtual reality paradigm

Participants were seated and observed an immersive first-person driving simulation from the perspective of a passenger in a vehicle moving forward automatically. On each trial an oncoming vehicle appeared in the opposite lane. Both conditions begin with the same event: the participant’s tire blows out, delivered as a loud bang together with the visible burst and a swerve of the vehicle. This tire blowout is the moment at which the two conditions diverge, and is the event to which the event-related potentials (ERPs) analyzed here are time-locked. On collision trials the oncoming vehicle then struck the participant’s vehicle; on no-collision trials the vehicle passed without incident (Figure 1). The impact itself was not written to the trigger channel, so its latency cannot be recovered from these recordings; it is bounded by the 3.49 s outcome phase that follows the blowout on collision trials (Trial timing, below). The paradigm was entirely passive; no motor response was required at any point. The oncoming vehicle also served as gaze support: it approaches from the far end of the scene throughout the 7 s approach phase and so acts as a slowly moving fixation target, the function a stationary fixation cross serves in conventional EEG experiments, without which free viewing of a driving scene would leave gaze and attention uncontrolled across the trial.

**Figure 1.**
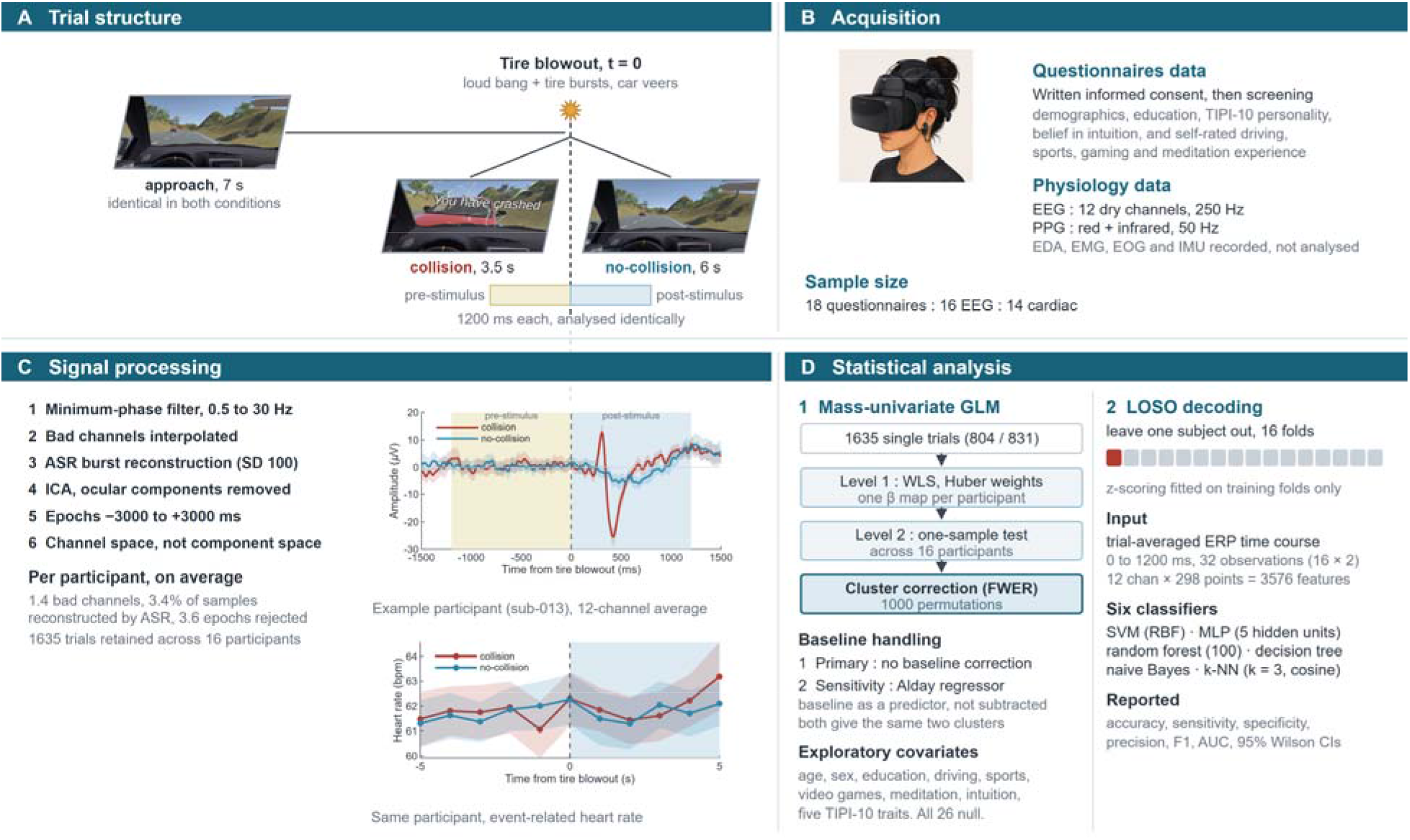
Study design, acquisition, signal processing and analysis. (A) Trial structure. A 7 s approach phase, identical in both conditions, ends in a tire blowout carrying both an auditory and a visual cue; the trial then resolves into a collision or no collision. Bars are drawn to scale from the measured median durations. The two analysis windows are symmetric about the blowout and are treated identically. Stills show the participant’s view in each condition. (B) Acquisition. (C) Signal processing, in order of application, with one example participant (sub-013) shown to illustrate the data the headset yields: ERPs averaged over the 12 channels and event-related heart rate, each with a 95% CI across that participant’s trials. These are single-participant illustrations, not results; the group analyses appear in Figures 3–5. (D) Statistical analysis: the two-level mass-univariate general linear model (GLM) with cluster correction, and the leave-one-subject-out (LOSO) decoding analysis.

The experiment used a within-participant repeated-measures design. Each participant completed 120 trials. Trial type was determined by a qRNG via the Australian National University Quantum Random Numbers Server API, with each trial assigned independently at 50% probability. Unique sequences were pre-generated per participant, stored as CSV files, and loaded automatically by the VR application at session onset. All sequences were validated blind, before data collection, by runs tests, autocorrelation, Shannon entropy and chi-square tests (Supplementary Material).

#### Trial timing

Timing was recovered from the device timestamps of the raw trigger channel for all 2,146 recorded trials. Every trial began with an approach phase in which the vehicle drove forward and the oncoming vehicle became visible; the tire-blowout marker, at which the conditions diverge, occurred 7.02 s after trial onset. This approach phase was effectively invariant and did not differ between conditions (collision 7.02 s, SD = 0.04; no-collision 7.03 s, SD = 0.06; d = −0.07), and its minimum across all trials was 6.49 s, so the entire analyzed epoch always fell within the current trial. The outcome phase that follows the blowout is shorter on collision trials, because the collision sequence ends a trial sooner: 3.49 s (SD = 0.06) against 6.01 s (SD = 0.04). The interval between successive blowouts was therefore not constant: 10.52 s following a collision and 13.03 s following a no-collision trial. That asymmetry depends on the preceding trial, which the quantum randomization makes independent of the current one, so the interval preceding a collision trial and preceding a no-collision trial did not differ (11.79 vs 11.77 s, d = 0.01). Temporal expectancy therefore cannot contribute to the condition contrast. Because sequences were generated by a hardware random source with equiprobable categories, the expectation-bias artifact described by Dalkvist et al. (2002) is not expected to operate in these data; the registered block analysis (Section 3.5) tests its signature directly.

### 2.4 Apparatus and acquisition

Physiological data were recorded with the Galea system (OpenBCI Inc.), a wireless multimodal headset integrating EEG, EDA, PPG, electromyography (EMG) and inertial measurement unit (IMU) sensors within a Varjo Aero VR head-mounted display (HMD). EEG was recorded from 12 scalp sites using dry pin electrodes in a modified 10-10 montage, in which two electromyography disc electrodes were reconfigured as EEG channels at Fp1 and Fp2. EEG was sampled at 250 Hz: the board reports that rate and no packets were dropped in any recording. The import routine used for the archived analyses derived 248 Hz from the median inter-sample interval, and every analysis reported here was run at that value, which scales reported latencies by 0.8% (at most 7 ms at the longest cluster, below the resolution the design supports). Because the sampling rate also governs filter coefficients and the segments retained by artifact subspace reconstruction (ASR), reproduction should start from the archived epoched datasets; the recovery procedure in Section 2.7 forces the archived rate for the same reason. PPG was recorded simultaneously from red and infrared channels at 50 Hz. EDA and eye-tracking sensors were present in the hardware but are not analyzed: no usable EDA signal was obtained in any recording, and the eye-tracking signal relies on analysis software that was still under development during data collection, so pupillometry was not available at the time the study was run (the preregistration update documents this in full).

Before the session, participants completed questionnaires covering eligibility, demographics, personality (TIPI), belief in intuition, and self-rated experience in driving, sports, video gaming and meditation. Participants were asked to wash their hair, avoid make-up and caffeine, and wear contact lenses rather than glasses where possible. Forehead and earlobe skin was cleaned with alcohol wipes, eye calibration was performed with the Varjo software, and electrodes were positioned as close to 10-10 nomenclature as the headset permitted, with pin electrodes rotated to penetrate the hair. Signal quality was verified with the manufacturer’s software before recording.

#### Reference

EEG was referenced to an ear-clip electrode on the earlobe, with the contralateral earlobe carrying the bias (ground) electrode, following the standard configuration for the OpenBCI ExG architecture on which the Galea is built. Signals were retained in this recording reference throughout; no re-referencing was applied at any stage. An average reference was not used: with 12 electrodes distributed unevenly over the scalp, the average of the recorded channels is a poor estimate of a neutral reference and introduces a spatial bias of its own, so the transformation would degrade rather than improve the data. The consequence is that the scalp distribution of an effect is tied to the recording reference and cannot be disentangled from it at this channel count. Peak electrodes are therefore reported as descriptive maxima, and no inference is drawn from the topography.

### 2.5 EEG preprocessing

Preprocessing was performed in MATLAB R2026a using EEGLAB 2026.0.0. Continuous data were trimmed to 1 s before the first and 1 s after the last event marker. A minimum-phase causal bandpass filter (0.5–30 Hz) was applied to preserve temporal causality and prevent post-stimulus activity from smearing into the pre-stimulus window (Widmann et al., 2015; Alday, 2019). A duplicate dataset was high-pass filtered at 1 Hz with the same filter type for independent component analysis (ICA).

Because the two prefrontal channels converted from disc electrodes were sometimes recorded with inverted amplifier leads, an automated polarity check was applied after filtering (sign agreement with a robust median reference at the top 10% of reference-amplitude time points, after an on-the-fly 8 Hz low-pass; channels below 0.5 agreement reversed).

Bad channels were flagged with a sliding-window procedure (2 s windows, 50% overlap) adapted from the clean_rawdata plugin. Within each window, the absolute correlation of each channel with all others was computed and the lowest 20% of correlations discarded; a channel was flagged in that window if it was both an amplitude outlier across windows and had all remaining correlations below 0.55, or if it was flat (maximum absolute first difference < 1e-7). Channels flagged in more than 30% of windows were removed. ASR (threshold = 100) was applied to the 1 Hz high-pass dataset, and the same time segments were removed from the 0.5 Hz dataset. Removed channels were interpolated by spherical spline before ICA.

ICA was performed with the Preconditioned ICA for Real Data algorithm (Picard; Ablin et al., 2018), taking the effective data rank into account (Kim et al., 2023). Decomposition weights were transferred to the 0.5 Hz dataset. Given the low channel count and the dry-electrode montage, automatic component classification was unreliable; the first component, which in this configuration consistently captured ocular activity, was removed after visual confirmation for every participant. One participant required removal of a second ocular component.

The full pipeline is implemented as an EEGLAB plugin released with this study (version 1.7.1, https://github.com/amisepa/galea-eeglab-plugin); its two dialogs, set as used for this study, are shown in Figure 2.

**Figure 2.**
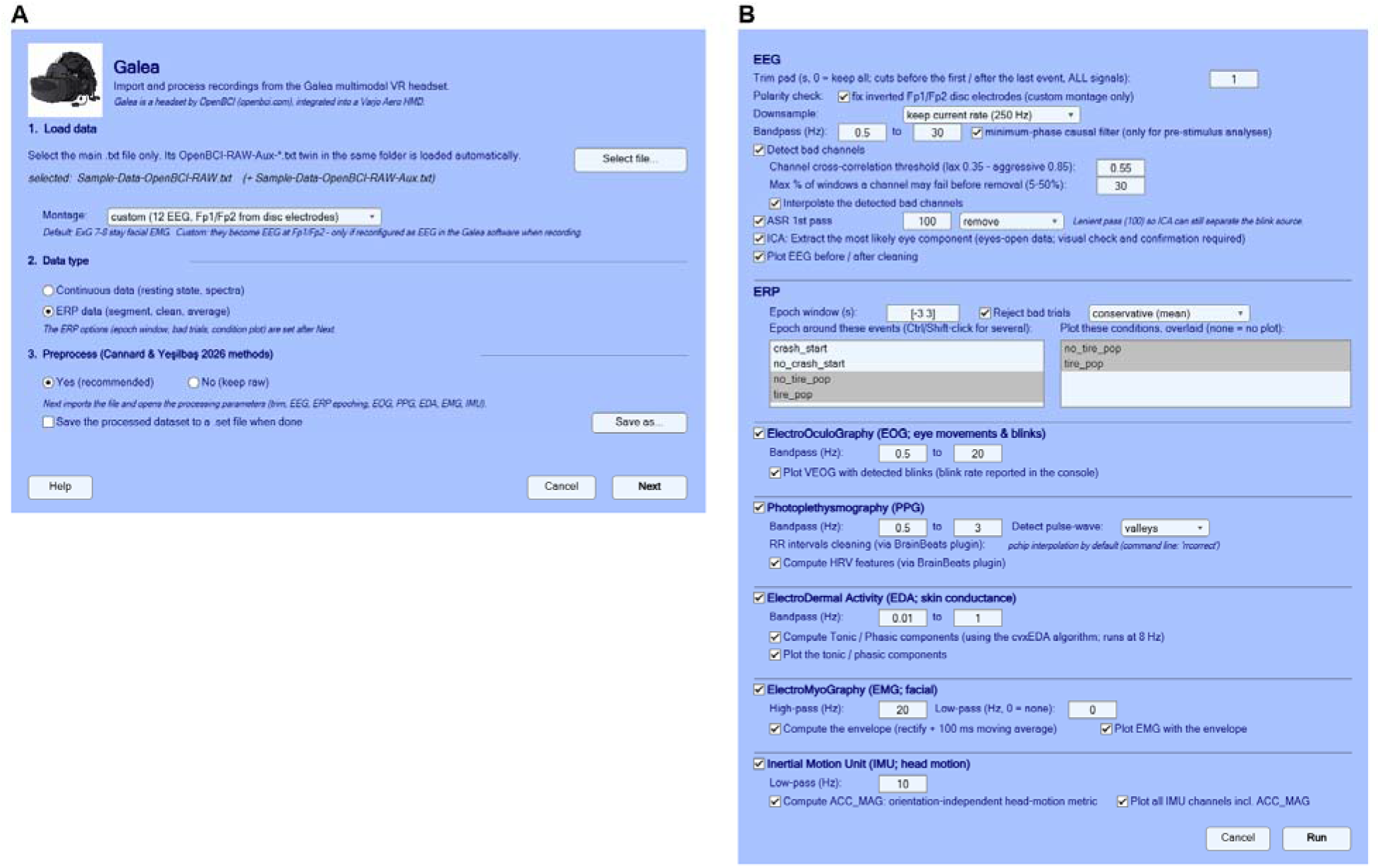
The EEGLAB plugin released with this study, set as used here. (A) Main window: the Galea/OpenBCI main file is selected (its Aux twin, carrying PPG, EDA and IMU, is found automatically), then the montage (custom: the two disc electrodes recorded as Fp1/Fp2), the data type (continuous or ERP) and whether to preprocess; Next imports the file and opens the processing parameters. (B) Processing parameters for ERP data, applied on Run: trimming to 1 s around the first and last event marker, 0.5–30 Hz minimum-phase causal bandpass, Fp1/Fp2 polarity check, bad-channel detection (correlation threshold 0.55, removal above 30% of windows) with interpolation, ASR at threshold 100 in remove mode (flagged segments deleted, with any event markers inside them reported), and ocular independent component removal with confirmation; ERP epoching from −3 to +3 s around the tire-pop and no-tire-pop events, with bad-trial rejection (mean-based criterion) and both conditions overlaid (20% trimmed mean with 95% confidence interval); and the electrooculography, PPG (cleaning of inter-beat (RR) intervals and heart rate variability features via the BrainBeats plugin), EDA, EMG and IMU branches.

Data were segmented from −3000 to +3000 ms relative to the tire-blowout markers (event codes tire_pop and no_tire_pop), that is, the onset of the divergence between conditions. Epochs with outlier root-mean-square amplitude or outlier high-frequency residual power were rejected using the mean-based criterion of MATLAB’s isoutlier function. No baseline correction was applied, in order to treat the pre- and post-stimulus periods identically and avoid introducing artificial differences between conditions (Maess et al., 2016; Delorme, 2023); normalization for time-frequency analyses is described in Section 2.7. Within-participant averages were inspected visually to confirm clear stimulus-locked deflections and adequate signal quality.

### 2.6 Time-domain statistical analysis

Mass-univariate GLMs were computed across all electrodes and time points, separately for the post-stimulus (0 to 1200 ms) and pre-stimulus (−1200 to 0 ms) periods, with trial type (collision versus no-collision) as the predictor. The two windows were deliberately matched in length so that any difference between them cannot be attributed to unequal numbers of comparisons or unequal cluster opportunity.

A two-level hierarchical approach accommodated the unbalanced and variable trial counts across participants. At the first level, a trial-wise weighted least-squares (WLS) GLM with Huber weights was fitted within each participant, down-weighting outlier trials without discarding them and yielding a participant-specific map of the collision-minus-no-collision effect. At the second level, these maps were submitted to a one-sample test across participants. Inference used a permutation null distribution constructed from 1000 iterations in which condition labels were shuffled within each participant at the first level before recomputing the entire two-level procedure, thereby preserving the within-participant trial structure. Family-wise error was controlled with cluster-based spatiotemporal correction across electrodes and time points at α = 0.05. Clusters separated by gaps shorter than 10 ms were merged before reporting. Effect sizes are Cohen’s d computed from the distribution of participant-level estimates.

As a sensitivity analysis, the same models were refitted with the per-trial baseline entered as an additional Level-1 predictor rather than subtracted (Alday, 2019), using a window of −3000 to −2000 ms common to both analysis windows. That window lies inside the approach phase of every trial, ends 800 ms before the pre-stimulus window begins, and is identical for both analyses, so it preserves the symmetry of the design. Entering the baseline as a regressor lets its weight be estimated from the data rather than assuming a coefficient of −1, and avoids injecting baseline noise into every trial estimate.

#### Cluster correction

Family-wise error in the time-domain analyses was controlled by cluster-mass permutation correction, implemented in analysis/cluster_correct.m. The time-frequency analyses use a different cluster statistic and are described in Section 2.7; the two are not interchangeable and we state which was used wherever a corrected result is reported. Clusters were formed at a two-tailed t threshold corresponding to p = 0.05, positive and negative excursions were clustered separately so that adjacent effects of opposite sign cannot merge, cluster mass was the sum of |t| within a cluster, and the observed and null maps were thresholded by the identical rule. The null distribution was the maximum cluster mass over 1000 permutations and corrected p-values include the observed statistic in the null, so no p-value can be exactly zero (Phipson and Smyth, 2010). Channel adjacency was defined by Delaunay triangulation of the electrode montage, giving a mean of 4.7 neighbors per channel. Clustering used standard adjacency, with no minimum-neighbor criterion. Such a criterion has no principled setting on a montage this sparse: adjacent electrodes are several centimeters apart, so a genuine effect need not be expressed at two of them, and requiring it would erode spatially broad and focal effects alike.

### 2.7 Time-frequency analysis

Because this study tests for a pre-stimulus difference, the time-frequency estimator was chosen so that it cannot move post-stimulus signal backwards in time. A conventional symmetric Morlet wavelet integrates over a window centered on the sample being estimated, so power at a pre-stimulus latency is partly determined by what happens after the stimulus; at the low frequencies of interest that window is hundreds of milliseconds wide, which is large relative to the effect being sought. Estimates were therefore computed with a causal (one-sided) Morlet wavelet, truncated at zero lag so that each sample is a function of preceding data only, and applied by forward-only convolution with left-side padding. Mirror padding was not used, as reflecting the signal about the trial edge reintroduces post-stimulus information into the pre-stimulus estimate.

Wavelets spanned 3 to 30 Hz in 0.5 Hz steps, with the number of cycles increasing linearly from 4 to 8 across that range. The lower bound reflects causality: a wavelet has temporal support of approximately ±3σ with σ = cycles / (2πf), so at 1 Hz it draws on nearly two seconds of data, and no pre-stimulus estimate within a second of stimulus onset can be kept free of post-stimulus signal. At 3 Hz the one-sided support is about 640 ms, which the analysis window accommodates. Instantaneous power was taken as the squared modulus of the convolution output and averaged across channels with a 20% trimmed mean. The time-course panel of Figure 4E was derived from the same trial-level power: per trial, in dB (10·log), averaged over the same channels and over trials within each participant and condition, then averaged across participants (mean ± 1 s.e.m.).

Two normalizations were applied in parallel. The primary analysis applies none: raw power enters the model directly, in decibels, because conventional baseline correction divides every estimate by mean power in a pre-stimulus window and so forces the corrected difference toward zero inside that window. The robustness check instead enters baseline power as a regressor rather than dividing by it (Alday, 2019), which adjusts for pre-trial power differences while leaving the pre-stimulus contrast free to vary. Baseline power was taken from −2300 to −1900 ms, separated from the analyzed pre-stimulus interval by more than the one-sided support of the lowest-frequency wavelet, so the covariate and the tested signal share no samples.

The identical hierarchical GLM and permutation procedure was then applied, with cluster correction operating over the joint frequency-time space. Time-frequency clusters were formed at an uncorrected threshold of p < 0.05 and assessed against the permutation distribution of maximum cluster extent, whereas the time-domain analysis used cluster mass; both control the family-wise error rate at alpha = 0.05. Two properties of the time-frequency correction differ from the time-domain one and are stated for completeness: the cluster statistic is the number of points in a cluster rather than the sum of |t| within it, and positive and negative excursions are thresholded on |t| and so could in principle merge. Neither is consequential for the results reported here, because every suprathreshold point in both windows is positive, so no cluster contains excursions of both signs. Pre-stimulus and post-stimulus windows were corrected separately, so no post-stimulus cluster can contribute to a pre-stimulus one.

Truncating a Gaussian-enveloped wavelet at zero lag leaves a kernel with spectral sidelobes that decay far more slowly than those of the symmetric wavelet it is derived from, so the effective bandwidth is broader than the nominal cycle count implies and adjacent frequency rows are strongly correlated. The analysis therefore resolves the presence and timing of spectral power changes but not their frequency specificity: peak frequencies are the center of a broad, correlated response rather than a band assignment, and an effect extending across the analyzed range is not by itself evidence that the underlying change is broadband. Amplitude normalization is unaffected (measured steady-state response flat to within 3% from 3 to 30 Hz), as is the causality guarantee, which follows from the truncation itself.

Pre-stimulus control analyses. Where a pre-stimulus cluster survived correction, the registered control analyses were run on mean power within it, one value per trial. Two of these require each analyzed epoch to be located in the delivered 120-trial sequence. The event markers in the raw recordings and the pre-generated quantum-random-number sequences identify every delivered trial’s position and condition; what the archived data do not store is which of the 120 trials survived preprocessing: ASR removes segments from the continuous recording and epochs spanning the resulting discontinuities are then dropped, so the retained set is a non-contiguous subset and the removed-segment mask was not saved. The mapping was recovered by replaying the length-determining steps of the pipeline (filtering, the stored bad-channel selection, and ASR) and reading off the surviving events; ICA was not replayed (it alters the data but not its length and creates no discontinuities). Each recovered mapping was verified against values stored by the original run (proportion of samples reconstructed, per-condition trial counts, epoch count); one recording did not reproduce and its participant is excluded from these analyses only, leaving 15. The same recovered index was used to align the ocular and EMG channels, which never underwent ASR and so have a different epoch set, trial by trial to the EEG.

Time-symmetry analysis. The time-symmetry correlation (hypothesis H4) is peak-matched to the primary mass-univariate effects, which here are the two time-frequency clusters (Section 3.4; Section 2.1). For each trial of every participant, the index is the mean channel-averaged power within the pre-stimulus cluster window and within the post-stimulus cluster window (the windows of Figure 4). Association was measured with the robust estimator the registration names (skipped Spearman; Pernet et al. (2013), Robust Correlation Toolbox), tested by 10,000 within-participant permutations of the pre–post pairing, and combined across participants with a one-sample t test on Fisher-transformed coefficients. Because the windows come from these data, the result is exploratory.

### 2.8 Classification analysis

To test whether condition could be decoded from held-out participants, ERPs were averaged across trials within each participant and condition, and the resulting waveform was used as the feature vector. Averaging is across trials only: the time course is preserved, so each observation is 12 channels × 298 samples = 3,576 features. The windows are the symmetric pair used by the GLMs, 0 to +1200 ms and −1200 to 0 ms, so that the mass-univariate and multivariate analyses interrogate the same interval. This gives 32 observations (16 participants × 2 conditions). Six classifiers were evaluated: a support vector machine with a radial basis function kernel, a multilayer perceptron with a single five-unit hidden layer, a Gaussian naive Bayes classifier, a random forest with 100 trees, a decision tree, and k-nearest neighbors (k = 3, cosine distance, inverse distance weighting). Evaluation used LOSO cross-validation, with both of a participant’s condition averages held out together and predictions pooled across folds before computing metrics. Features were z-scored using means and standard deviations computed from training folds only. The identical procedure was applied to the pre-stimulus window. Given the small number of observations, accuracies are accompanied by 95% Wilson score intervals. Significance was assessed against a permutation null: condition labels were shuffled within participant, preserving each participant’s pair structure, and the entire cross-validation was repeated 1000 times to build the null distribution of accuracy. This matches the permutation scheme used by the mass-univariate analyses, so the two families of tests rest on the same exchangeability assumption. The smallest attainable p-value is 1/1001.

The same procedure was then applied to the cardiac data and to the two modalities combined. The cardiac feature vector is the participant’s mean instantaneous heart rate sampled at eleven points spanning the full −5 to +5 s epoch, used alone and concatenated onto the ERP features. Note that this is a wider interval than the ±1200 ms used for EEG: heart rate resolves at roughly one sample per beat, so a 1.2 s window would contain one or two values and could not support a classifier at all. The three feature sets are therefore matched on participants and folds but not on time window, and the comparison should be read accordingly. Because the cardiac quality screen retained a different subset of participants from the EEG screen, this three-way comparison is restricted to the 14 participants with usable data in both modalities, so that all three feature sets are evaluated on identical folds and are directly comparable (28 observations). The concatenation is reported for completeness rather than as a competitive fusion: appending eleven features to several thousand cannot be expected to move a decision boundary, and it is the heart-rate-only model that establishes whether cardiac information is present at all.

### 2.9 Cardiac data

PPG signals were trimmed to the experimental period with a 1 s buffer. Signal quality was assessed per channel with a frequency-domain metric: each channel was high-pass filtered at 0.75 Hz and segmented into 4 s windows with 50% overlap, and signal-to-noise ratio was computed for each window as the decibel ratio of power within cardiac bands (0.8–2.5, 1.6–5.0 and 2.4–7.5 Hz, capturing the fundamental pulse rate and its harmonics) to power outside them, with the median across windows as the channel index. The infrared channel was superior in every retained participant (signal-to-noise ratio 3.9–19.2 dB) and was selected throughout. It was then bandpass filtered between 0.5 and 3 Hz with a causal minimum-phase FIR filter.

Four participants were excluded: three for insufficient signal quality and one whose session was interrupted and re-started, leaving N = 14. Heartbeats were detected and converted to RR intervals with the BrainBeats plugin (Cannard et al., 2024). Artifactual and ectopic beats were removed by an automated procedure, flagging a mean of 0.7% of beats (SD = 0.8, range 0.0– 2.8). Instantaneous heart rate was derived from the normal-to-normal (NN) beat intervals as 60/NN and epoched from −5 to +5 s around each event, without baseline correction. No-collision markers within 5 s of one another were de-duplicated and any coinciding with a collision event removed. Trials with outlier root-mean-square values were excluded per participant, retaining a mean of 58.1 collision trials (SD = 4.2, range 52–66) and 61.3 no-collision trials (SD = 7.6, range 39–69).

Condition differences were assessed with the same two-level hierarchical WLS/Huber GLM and permutation procedure described in Section 2.6, applied separately to the post-stimulus (0 to +5 s) and pre-stimulus (−5 to 0 s) windows, with multiple comparisons across time points controlled by t-max correction at α = 0.05. Confidence intervals on the group differences were constructed with a percentile bootstrap (2000 resamples over participants).

### 2.10 Exploratory individual-difference analyses

Thirteen individual-difference variables (age, sex at birth, years of education, self-rated driving, sports, video-game and meditation experience, belief in intuition, and the five TIPI traits) were each entered separately as a second-level covariate in the hierarchical GLM, in both time windows: 26 models in all (13 variables × 2 windows). For the cardiac data, the per-participant post- and pre-stimulus heart-rate condition effect was correlated with the same variables using skipped Spearman correlations (Pernet et al., 2013), with bivariate outliers identified by projection onto the minimum covariance determinant center and a chi-square-based box-plot rule, followed by Benjamini–Hochberg false discovery rate (FDR) correction across the 13 variables, as registered.

For the EEG models the registered FDR correction across variables is non-binding. Each of the 26 was cluster-mass corrected within itself and none produced a surviving cluster. Benjamini– Hochberg is a step-up procedure whose adjusted p-values are never smaller than the raw ones, so a family containing no rejection before correction cannot contain one after it; with no rejections the FDR across these models is zero by construction. The individual-difference results are reported as exploratory and presented in Supplementary Material rather than as findings of this study on grounds of statistical power, not because multiplicity was left uncontrolled.

## 3. Results

### 3.1 Data quality

Across the 16 retained participants, a mean of 1.44 channels per participant (SD = 1.15, range 0– 3) were flagged as bad and interpolated. ASR removed a mean of 3.36% of continuous data (SD = 3.22, range 0.03–11.89). A mean of 3.56 epochs per participant (SD = 0.96, range 2–6) were subsequently rejected, leaving 50.3 collision trials (SD = 4.5, range 42–56) and 51.9 no-collision trials (SD = 9.8, range 35–64), for a total of 1635 trials entering the group analysis. The preregistered artifact-free trial minimum was relaxed before analysis for the four participants who fall below it in the no-collision condition, as documented in the preregistration update. The greater variability in no-collision counts reflects the independent 50% assignment of each trial by the qRNG.

### 3.2 Post-stimulus period: time domain

The hierarchical GLM revealed two significant clusters distinguishing collision from no-collision trials in the post-stimulus window (Figure 3; Table 1). Cluster 1 (negative) extended from 363 to 665 ms, peaking at 452 ms over C3 (t = −5.39, d = −1.35, p = .001), and involved 11 electrodes; amplitudes were more negative on collision than on no-collision trials. Cluster 2 (positive) extended from 778 to 1198 ms, peaking at 1028 ms over O1 (t = 3.85, d = 0.96, p = .001), and involved 8 electrodes; amplitudes were more positive on collision than on no-collision trials.

**Figure 3.**
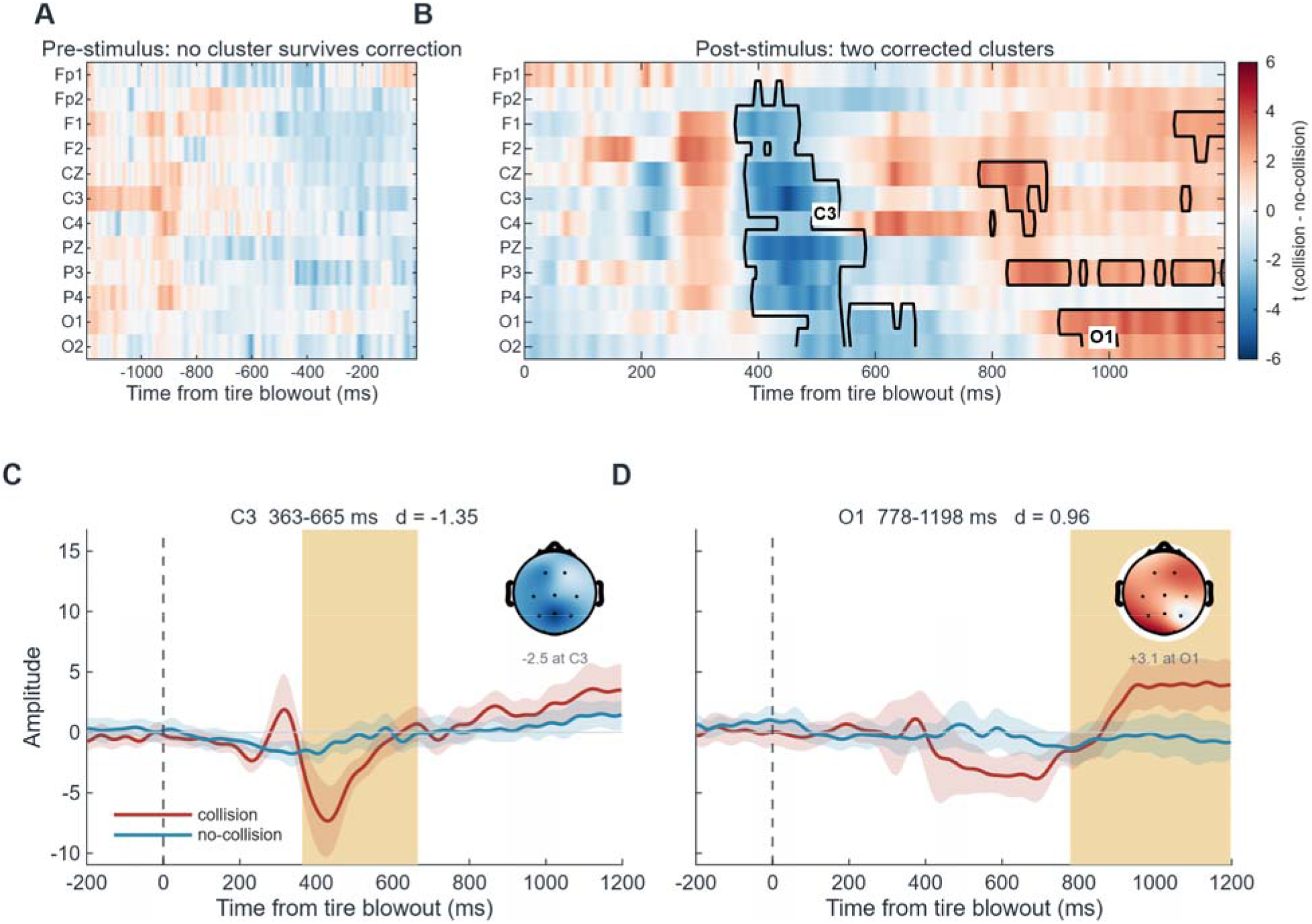
Mass-univariate results in the time domain, collision minus no-collision. (A, B) t-values at every channel and time point in the pre- and post-stimulus windows, on a common color scale; black outlines mark the clusters surviving spatiotemporal permutation correction (alpha = 0.05, 1000 permutations, N = 16). No cluster forms in the pre-stimulus window. (C, D) Grand-average ERPs at the peak channel of each post-stimulus cluster, with 95% CIs across participants and the cluster extent shaded; insets show the mean difference topography over the cluster window. Traces carry a display-only 15 Hz low-pass; all statistics were computed on the analyzed 0.5–30 Hz data.

**Table 1.** Significant post-stimulus ERP clusters for the collision versus no-collision contrast, after cluster-based spatiotemporal permutation correction (alpha = 0.05, 1000 permutations, N = 16). Peak site is the electrode carrying the maximum absolute t-value within the cluster; d is Cohen’s d computed from participant-level estimates.

| **Cluster** | **Onset (ms)** | **Offset (ms)** | **Peak (ms)** | **Peak site** | **Electrodes** | **t** | **d** | **p (corr.)** |
|---|---|---|---|---|---|---|---|---|
| 1 | 363 | 665 | 452 | C3 | 11 | -5.39 | -1.35 | .001 |
| 2 | 778 | 1198 | 1028 | O1 | 8 | 3.85 | 0.96 | .001 |

All clusters survived cluster-mass permutation correction (α = 0.05, 1000 permutations). Peak sites are given as descriptive labels for where each effect was largest, not as claims about generators: 12 electrodes in a fixed hardware reference cannot localize, and an average reference is not available as a remedy at this channel count (Section 2.4). The interpretation rests on the timing and the magnitude of the effects, not on where they appear largest.

No cluster falls in the window predicted by hypothesis H1, which anticipated an early sensory response within approximately 100–150 ms of onset: nothing survives correction anywhere before 363 ms. The second part of that hypothesis, which predicted temporally extended differences over roughly 300–1000 ms, is consistent with the earlier of the effects reported above.

#### Sensitivity to baseline treatment

Refitting with the per-trial baseline as a Level-1 predictor rather than subtracting it (Alday, 2019) reproduced the same two clusters with essentially unchanged timing and effect sizes: 367–710 ms peaking at C3 (t = −5.21, d = −1.30), 774–1198 ms peaking at O1 (t = 3.86, d = 0.96). The baseline window itself showed no meaningful condition difference at any electrode (maximum |d| = 0.43, against 1.35 for the post-stimulus effect), so the regressor acts as variance reduction rather than competing with the condition term. The pre-stimulus window remained empty under this model as well.

### 3.3 Pre-stimulus period: time domain

No cluster survived correction in the pre-stimulus window. Only 2.5% of time-electrode points reached the cluster-forming threshold, against the 5% expected by chance, and the largest absolute t-value was 3.10. Clusters did form (26 of them), but the largest fell far short of the permutation threshold, which the post-stimulus window exceeded twice over. For comparison, 22.5% of points were suprathreshold post-stimulus. This window was identical in length to the post-stimulus window and was analyzed with the identical pipeline, so the contrast between the two periods cannot be attributed to differences in the number of comparisons, in cluster opportunity, or in analytic treatment. Because the data were filtered causally, the pre-stimulus signal cannot contain information leaked backwards from the post-stimulus response.

### 3.4 Time-frequency domain

After the primary analysis, which applies no normalization, the post-stimulus window contained one significant cluster in the joint frequency-time space (Figure 4). Cluster 1 (power increase for collision relative to no-collision) spanned 3.0–30.0 Hz and 266–766 ms, peaking at 7.0 Hz / 444 ms (t = 7.89, d = 1.97).

**Figure 4.**
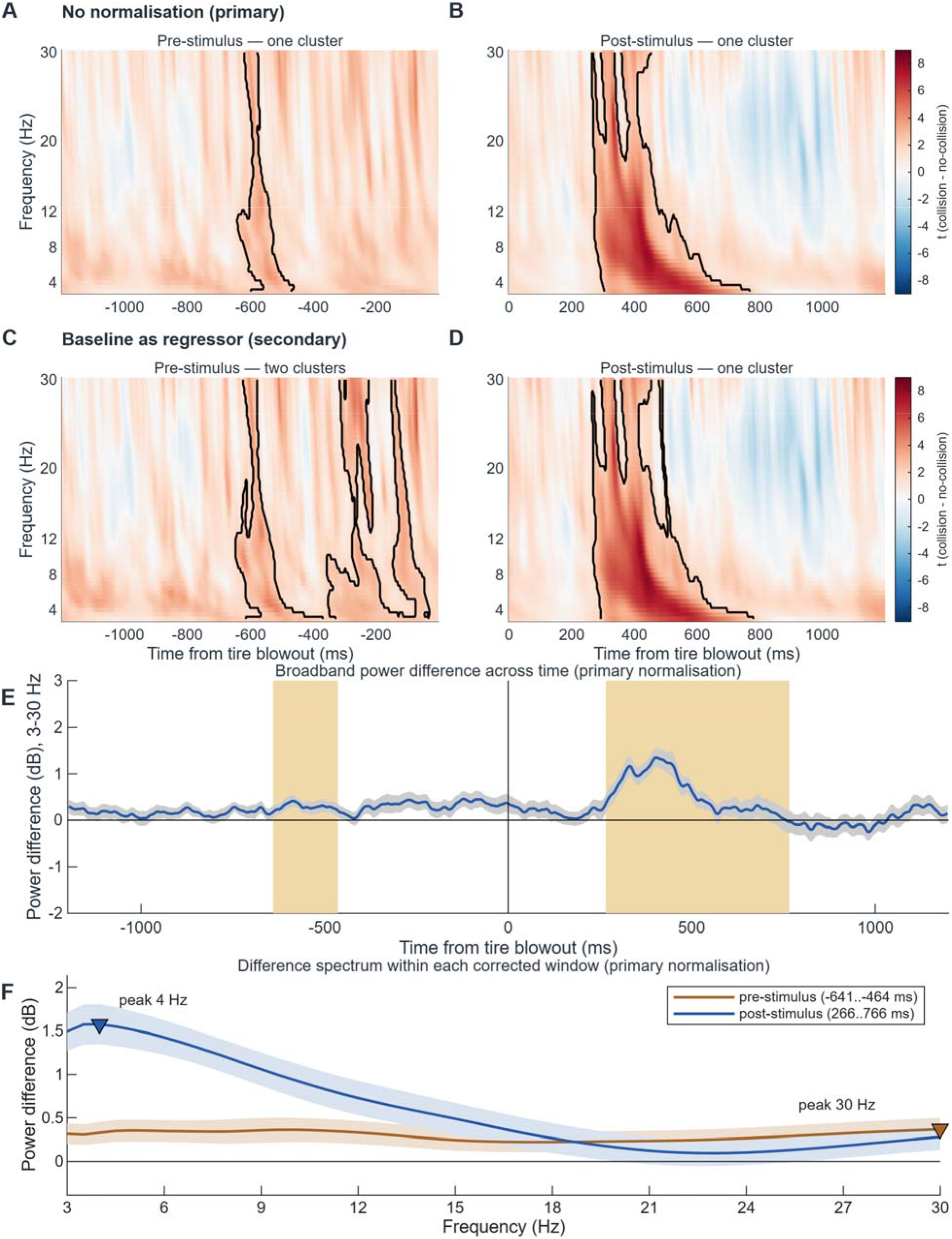
Time-frequency results, collision minus no-collision, averaged across channels, from the causal wavelet estimator, shown under both normalizations on a shared color scale. (A, B) No normalization, the primary analysis, pre- and post-stimulus. (C, D) Baseline power entered as a regressor (Alday, 2019), the secondary analysis. Black outlines mark clusters surviving cluster-extent permutation correction, corrected separately per window, and labels give the peak frequency of each; as noted in Section 2.7, peak frequencies are the center of a broad and correlated response rather than a band assignment. Note that the outlines mark where the effect crosses the cluster-forming threshold, not its extent: the pre-stimulus difference is positive across almost the whole panel (Section 3.5). (E) The same difference as a time course: per-trial power in dB, averaged over channels and trials per participant and then across participants (mean, solid line; ±1 s.e.m., band), with the corrected cluster windows shaded; it shows the same pattern as the maps, a small sustained pre-stimulus elevation and a larger post-stimulus peak. The two normalizations give the same answer: the pre-stimulus elevation is present under both, and the secondary analysis additionally resolves it into two sub-clusters (Section 3.4). (F) The difference spectrum within each corrected window, from the same trial-level power as (E), averaged over the window in time only (mean, solid line; ±1 s.e.m., band). The post-stimulus difference declines with frequency from 1.6 dB at 4 Hz, the shape of a broadband 1/f-like change; the pre-stimulus difference is comparatively flat (0.2–0.4 dB across 3–30 Hz), so the pre-stimulus effect is not driven by any one band, nor by the low frequencies that dominate the post-stimulus effect. Markers show the frequency of maximum difference (30 Hz pre-stimulus, 4 Hz post-stimulus); both sit at or near an edge of the analyzed range, which itself indicates that neither window contains a spectrally focused effect.

The baseline-as-regressor (Alday, 2019) analysis produced a closely comparable pattern: Cluster 1 (power increase for collision relative to no-collision) spanned 3.0–30.0 Hz and 266–778 ms, peaking at 7.5 Hz / 440 ms (t = 8.16, d = 2.04). The convergence between the two normalizations indicates that the post-stimulus oscillatory modulation is not an artifact of either choice.

In the pre-stimulus window the same analysis yielded: Cluster 1 (power increase for collision relative to no-collision) spanned 3.0–30.0 Hz and −641–-464 ms, peaking at 5.0 Hz / −540 ms (t = 3.98, d = 0.99). Given the exploratory status of this study, this requires independent replication. The control analyses in Section 3.5 characterize it and test the preregistered explanations for it.

Under baseline-as-regressor (Alday, 2019) the pre-stimulus window yielded two clusters (Figure 4C, D): Cluster 1 (power increase for collision relative to no-collision) spanned 3.0–30.0 Hz and −645–-371 ms, peaking at 5.0 Hz / −540 ms (t = 4.42, d = 1.11). Cluster 2 (power increase for collision relative to no-collision) spanned 3.0–30.0 Hz and −355–-32 ms, peaking at 26.0 Hz / - 246 ms (t = 4.83, d = 1.21). The effect is therefore present under both normalizations, and is more extensive under this one. All control analyses reported below were computed on the primary (unnormalized) window only.

### 3.5 Pre-stimulus control analyses

Because a pre-stimulus difference is the claim most vulnerable to mundane explanation, the cluster was examined further. All analyses in this section use mean power inside the corrected cluster, one value per trial. The run-length, block and peripheral analyses require each analyzed epoch to be located in the delivered 120-trial sequence and are therefore restricted to the 15 participants whose trial-to-sequence mapping could be recovered and verified (Section 2.7); the time-symmetry correlation requires no such mapping and uses all 16. Absolute p-values here are not independent evidence that the effect exists, because the window was selected for being extreme; what is interpretable is whether each control changes the effect, since the selection applies equally to both sides of every comparison.

The shape of the effect matters for how it should be read, and it is not the shape a cluster listing implies. The corrected cluster occupies 4.6% of the pre-stimulus frequency-time plane, but the difference is in the same direction across 94.0% of that plane, and of the 3284 points exceeding the cluster-forming threshold, every one is positive. Averaged in 100 ms bins the t-statistic does not fall below +0.7 at any latency in the window, and it is positive at every frequency from 3 to 30 Hz. The cluster boundaries therefore mark where a sustained offset happens to cross threshold, not where the effect begins and ends. We report the cluster because it is what the correction procedure returns, but the underlying difference is a broad elevation spanning the analyzed window rather than a transient at a particular latency, and the peak latency should not be interpreted as a moment of anticipation.

The effect is distributed across participants rather than carried by a few. It is in the same direction in 13 of 15 participants (sign test p = 0.007), with a group mean of +0.49 dB, and removing the single most extreme participant leaves it essentially unchanged at +0.42 dB.

Gambler’s fallacy and expectation bias (registered). With run length entered as a categorical predictor alongside condition, the condition effect is +0.53 dB, t(14) = 4.22, p < 0.001, essentially identical to its unadjusted value, and the alternative reading of the registered run-length definition gives the same answer (+0.52 dB, p < 0.001). Run length itself did not modulate pre-stimulus power: a partial F-test on the run-length block was significant in 0 of 15 participants (Fisher combination p = 0.960). A linear trend across run lengths was small and did not replicate across the two readings (−0.13 vs −0.02 dB per level), so we do not interpret it. In the registered complementary contrast, the condition effect was present after both long and short runs (+0.95 and +0.37 dB) and did not differ between them (+0.58 dB, t(14) = 1.33, p = 0.203).

CNV and time on task (registered). Trials were divided into early, middle and late blocks, excluding two participants for whom block membership is undefined (one whose session restarted after a headset disconnection, one whose recording begins at trial 30). The effect was +0.52 dB early, +0.27 dB in the middle and +0.77 dB late, with no increase from early to late (+0.24 dB, t(12) = 0.72, p = 0.485). The registration specifies that CNV and learning accounts predict a monotonic increase across blocks; that prediction is not met.

Ocular and muscular activity. Eye and facial-muscle activity volume-conducts to scalp electrodes, and a broadband pre-stimulus increase is what a blink or a jaw movement looks like, so the ocular and EMG channels recorded by the headset were analyzed on the same trials. Trial- to-trial coupling between peripheral and scalp broadband power is substantial and positive in 15 of 15 participants (mean partial r = 0.39 with condition removed), confirming that the two are not independent. However, peripheral power did not itself differ by condition (+0.34 dB, p = 0.188), and entering it as a covariate removed only 26% of the scalp effect, which remained at +0.34 dB (p = 0.002); the indirect path was not significant (p = 0.113). This is the expected pattern if peripheral and scalp signals share variance for reasons unrelated to condition. It does not exclude a peripheral contribution, which at this sample size cannot be estimated precisely, but the condition difference is not reducible to one.

Individual differences. The registered moderator hypothesis (H5) predicted that pre-stimulus differentiation would be stronger in participants with more experience in domains demanding sustained attention. The 13 questionnaire variables were correlated with each participant’s cluster-mean effect in both windows (26 tests, 13 of them pre-stimulus), with Benjamini– Hochberg applied across the family as registered. Nothing survives (smallest q = 0.81), and no association is close. H5 is not supported. With 14 to 16 participants this is a failure to reject rather than evidence of absence: the study was never powered for between-participant moderators, and the registration anticipated as much. All correlations are listed in Supplementary Table S2b.

Time symmetry (registered hypothesis H4, run post hoc). Time-symmetric accounts predict that trial-wise pre-stimulus differentiation and post-stimulus responses are two expressions of a shared process, and therefore covary. They do. For every participant, the correlation between trial-wise mean power in the pre-stimulus cluster window and in the post-stimulus cluster window is positive (16 of 16 positive; skipped Spearman, 10,000 pairing permutations per participant); the group-level 20%-trimmed mean coefficient is +0.29 (bootstrap 95% CI +0.25 to +0.39), and a one-sample t test on Fisher-transformed coefficients gives t(15) = 6.27, p < 0.001. The correlation survives when conditions are analyzed separately, so it is not merely shared condition variance. The windows come from these data, so the result is exploratory; per-participant values are listed in Supplementary Table S4.

No control removes the effect, and the registered dissociations point away from the explanations they were written to test; we nonetheless regard this as a finding requiring independent replication, for the reasons set out in Section 4.2.

### 3.6 Classification

Under LOSO cross-validation, all six classifiers separated the conditions from the post-stimulus response, with accuracies from 75.0% to 93.8% and areas under the curve (AUC) from 0.83 to 0.93 (Table 2). The best model reached 93.8% accuracy, 95% Wilson CI [79.9, 98.3]. Because performance is estimated from 32 observations the intervals are wide and the ordering of classifiers should not be over-interpreted; the informative result is that every model separated the conditions well above chance on participants it had never seen (all p ≤ .007, Table 2).

**Table 2.** Classification of collision versus no-collision from participant-averaged post-stimulus (0 to +1200 ms) ERPs, under LOSO cross-validation with predictions pooled across folds (16 participants, 32 observations). All values are percentages except AUC. Chance accuracy is 50%. p is the two-tailed permutation p-value against a within-participant label-shuffling null with 1000 permutations; the smallest attainable value is 1/1001.

| **Classifier** | **Accuracy** | **Sensitivity** | **Specificity** | **Precision** | **F1** | **AUC** | **p** |
|---|---|---|---|---|---|---|---|
| SVM | 78.1 | 81.3 | 75.0 | 76.5 | 78.8 | 0.87 | <.002 |
| k-nearest neighbours | 81.3 | 75.0 | 87.5 | 85.7 | 80.0 | 0.93 | <.002 |
| Naive Bayes | 75.0 | 75.0 | 75.0 | 75.0 | 75.0 | 0.83 | 0.007 |
| Random forest | 81.3 | 68.8 | 93.8 | 91.7 | 78.6 | 0.90 | 0.003 |
| Decision tree | 93.8 | 87.5 | 100.0 | 100.0 | 93.3 | 0.88 | <.002 |
| Neural network (MLP) | 84.4 | 75.0 | 93.8 | 92.3 | 82.8 | 0.93 | <.002 |

Applied to the pre-stimulus window with everything else held identical, the same procedure left every classifier at chance (Table 3). Accuracy ranged from 31.3% to 50.0%, and even the best model’s interval, [33.6, 66.4], spans 50%. All six AUCs fall slightly below chance (0.34–0.42); at 32 observations this is within sampling noise for a null effect and should not be read as inverted information. No classifier approaches significance against the permutation null (all p ≥ .57). Hypothesis H6 is not supported.

**Table 3.** The identical analysis applied to the pre-stimulus window (−1200 to 0 ms). Cross-validation scheme, feature construction and classifiers are as in Table 2; only the window differs.

| **Classifier** | **Accuracy** | **Sensitivity** | **Specificity** | **Precision** | **F1** | **AUC** | **p** |
|---|---|---|---|---|---|---|---|
| SVM | 43.8 | 18.8 | 68.8 | 37.5 | 25.0 | 0.37 | 0.808 |
| k-nearest neighbours | 31.3 | 43.8 | 18.8 | 35.0 | 38.9 | 0.35 | 0.959 |
| Naive Bayes | 50.0 | 68.8 | 31.3 | 50.0 | 57.9 | 0.42 | 0.574 |
| Random forest | 43.8 | 31.3 | 56.3 | 41.7 | 35.7 | 0.36 | 0.775 |
| Decision tree | 40.6 | 37.5 | 43.8 | 40.0 | 38.7 | 0.34 | 0.736 |
| Neural network (MLP) | 34.4 | 50.0 | 18.8 | 38.1 | 43.2 | 0.38 | 0.942 |

This agrees with the mass-univariate time-domain analysis, which also found nothing before onset. A multivariate decoder can in principle detect distributed patterns that a channel-wise GLM misses, so its failure here indicates that the evoked response carries little or no discriminative information in the pre-stimulus window, rather than that such information is merely undetectable by univariate methods. This does not extend to the spectral difference reported in Section 3.5. The classifier was given the event-related waveform, not time-frequency power, so it was never in a position to detect that effect and its null says nothing about it.

Finally, the two modalities were combined on the 14 participants with usable data in both. Heart rate alone was at chance (46.4–53.6% accuracy, AUC 0.46–0.53), which agrees with the univariate cardiac result reported below. Concatenating the heart-rate features onto the ERP moved mean accuracy from 82.1% to 81.5%, and four of the six classifiers returned predictions identical to the EEG-only model on every observation. Taken alone that null would be uninformative: eleven cardiac features appended to 3,576 electroencephalographic ones cannot be expected to shift a decision boundary whether or not they carry signal. The heart-rate-only result is what settles the question, and it indicates there was nothing in the cardiac response for the combination to add. Full metrics for all three feature sets are given in Supplementary Table S3.

### 3.7 Cardiac response

The hierarchical permutation GLM revealed no significant difference in event-related heart rate between conditions in the post-stimulus window (0 to +5 s; all permutation p > 0.37; Figure 5). No significant anticipatory difference was observed in the pre-stimulus window (−5 to 0 s) after correction, although a single time point near −4 s reached a nominally negative value (t ≈ −2.3, uncorrected p = 0.033) that did not survive correction. Both conditions produced closely overlapping absolute heart-rate trajectories averaging approximately 71–72 bpm, with wide CIs reflecting substantial between-participant variability (Figure 5).

**Figure 5.**
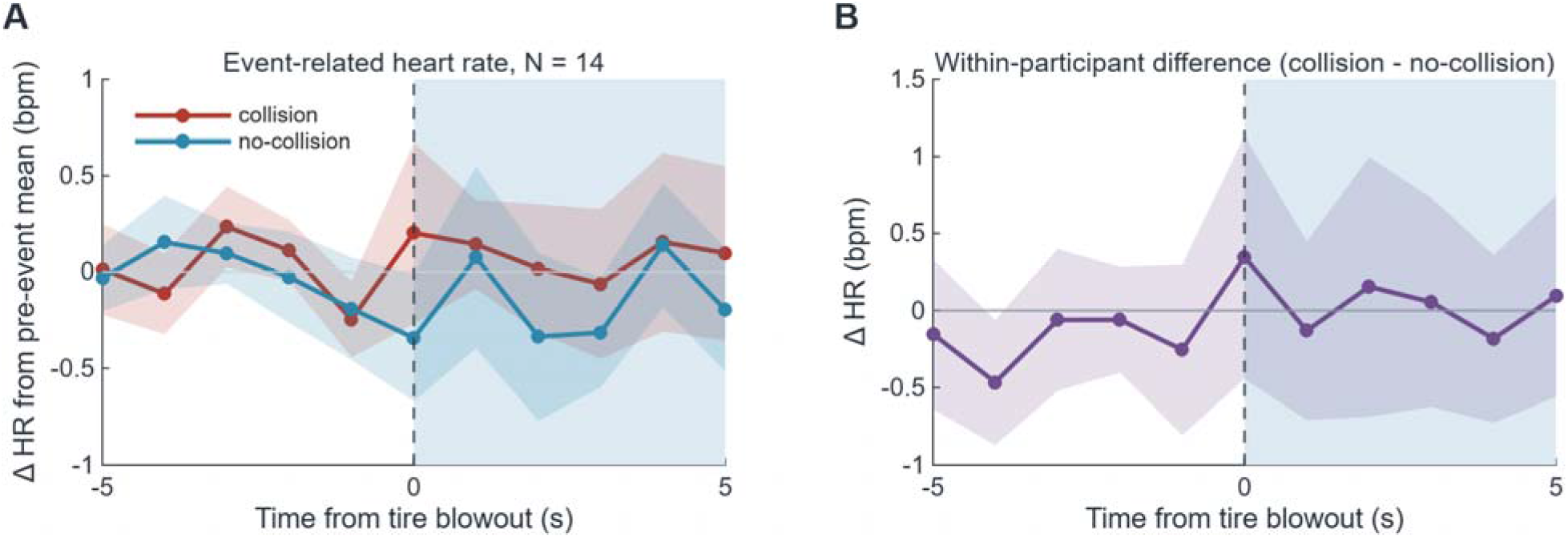
Event-related heart rate (N = 14). (A) Each condition referenced to its own pre-event mean, with 95% CIs across participants; absolute rate differs by tens of beats per minute between people, so the raw traces are dominated by between-participant spread. (B) The within-participant difference, which is the quantity tested. No time point survives t-max correction across the 11 time points in either window; the nominal dip near −4 s is uncorrected.

### 3.8 Exploratory individual differences

None of the 26 individual-difference models in the time domain produced a cluster surviving correction, in either window. This held for every demographic, experiential and personality variable tested, and is true even before any correction across the family of models is considered. Personality variables were available for 14 of the 16 participants and all other variables for all 16.

In the cardiac data, no variable survived FDR correction in either window (all corrected p > 0.05), although nominally significant associations appeared consistently across both windows for extraversion, years of education and emotional stability. Given the number of models and the sample size, none is interpreted here.

## 4. Discussion

We asked when discriminative neural information about an unpredictable collision becomes available, using an immersive VR driving paradigm designed to remove the confounds that complicate interpretation of earlier work. With causal filtering, no baseline correction, and pre- and post-stimulus windows matched in length and analyzed identically, evoked and decodable differentiation between collision and no-collision trials was found after stimulus onset, while a difference in broadband spectral power was present before it.

### 4.1 Post-stimulus differentiation

The post-stimulus response comprised two spatiotemporal clusters, the larger spanning 363–665 ms and peaking near 452 ms (d = −1.35). The corrected statistics show that the conditions diverge from roughly 363 ms after the tire blowout, that the divergence persists for several hundred milliseconds in the window in which evaluative processing of salient events is typically observed, and that it is accompanied by a broadband increase in spectral power over the same interval. The 12-channel montage cannot support named components or scalp localization, and we make no claim about either. This is the expected signature of a multisensory, survival-relevant event being detected and evaluated, and its magnitude is substantially larger than the effects typically reported in screen-based paradigms.

A second, later cluster spanned 778–1198 ms (d = 0.96), running to the end of the analyzed window. A sustained late difference of this kind is what would be expected if the collision continued to be processed after the initial evaluative response. It has no time-frequency counterpart: the post-stimulus spectral cluster overlaps the earlier time-domain cluster and resolves before this one begins. It is reported with more caution because the analyzed window ends at 1200 ms, so its resolution cannot be established.

These responses were resolved with dry pin electrodes, through hair, inside a VR headset, in participants whose mean age was 57 years, and they survived conservative cluster correction with fewer than 20 participants. Wireless multimodal systems are often proposed for applied neuroergonomics but are seldom validated against a demanding electrophysiological criterion. These data provide that validation for event-related responses to naturalistic threat, and the decoder that generalizes across participants indicates that the signal is consistent enough in form to be recognized in individuals the model has not seen.

### 4.2 Pre-stimulus period

The pre-stimulus result is mixed. No effect appeared in the time domain, and a classifier trained on the pre-stimulus window performed at chance on held-out participants. In the time-frequency domain, a broadband cluster survived correction several hundred milliseconds before an event whose outcome had not been determined by anything the participant could observe. This comparison was preregistered as the primary anticipatory hypothesis, and we do not regard the outcome as established.

The explanations available for it were preregistered, and none accounts for it (Section 3.5). Trial type was assigned by a quantum random source at equiprobable rates, and the registered run-length control confirms that the effect does not depend on the preceding sequence. Temporal expectancy was controlled by design rather than by jitter: varying the inter-stimulus interval randomly, a common control, does not remove expectancy, because the brain learns the interval distribution itself and times its preparation to the most probable onsets (Mento et al., 2013). Here the approach phase was fixed at 7.02 s in both conditions, so when the event would occur was predictable and only whether it would be a collision was not. The registered block analysis was designed to separate a CNV or learning effect from a predictive one by whether the effect grows across the session, and it does not grow. Minimum-phase causal filtering and causal wavelets prevent post-stimulus samples from entering the pre-stimulus estimate, and the windows were matched in length and corrected separately. The effect is present in 13 of 15 participants and survives removal of the largest contributor, so it is not an artifact of one or two outliers.

Three considerations limit interpretation. This is an exploratory analysis of a study stopped at 18 of 63 planned participants, and the estimate comes from a window chosen because it was extreme, so its magnitude is inflated by an unknown amount. It appears in one domain and not the other two. Ocular and muscular power did not differ by condition, but those channels are strongly coupled to the scalp signal trial by trial, so a peripheral contribution cannot be excluded at this sample size.

The form of the effect also constrains interpretation. It is not a transient rising toward onset but a sustained elevation across the analyzed window and band (Section 3.5). A transient would suit a stimulus-specific anticipatory process; a sustained offset is also what a tonic state difference between the two sets of trials would produce, and if collision trials were drawn from periods of systematically higher broadband power, the contrast would look like this. Under this design such a displacement has no ordinary source, because outcome was assigned by a quantum random source independently of everything preceding the trial, and trial loss was condition-symmetric. We record the shape because it bears on which explanations remain open, not because we can adjudicate between them.

If anticipatory activity is real but weak in stripped-down settings, an immersive paradigm is where a stronger version should appear, and this study was designed as that first probe. It ended far short of its planned sample and cannot settle the question, but the pre-stimulus difference is, to our knowledge, the first reported inside an immersive, ecologically valid environment. Its presence here, against its absence in the time domain and the classifier, is the pattern most worth resolving in an adequately powered study.

We therefore make *no claim* that pre-stimulus differentiation occurred, and no claim that it did not: the cluster survived its own preregistered controls, and omitting it would misrepresent the data. It requires independent replication in an adequately powered sample before it can be interpreted.

#### A caveat on the spectral findings

The post-stimulus time-frequency effect is a broadband power increase, not a modulation confined to a band. A uniform increase of this kind is what a shift in the 1/f-like background would produce, and Gyurkovics et al. (2022) show that such shifts follow stimulus onset and violate the stationarity assumption behind baseline normalization. We cannot distinguish that from genuine oscillatory modulation, and as Section 2.7 notes, the estimator cannot resolve the frequency axis finely enough to try. Neither normalization used here separates a time-varying aperiodic rotation from oscillatory modulation, and neither was designed to: the primary analysis applies no normalization at all, and entering baseline power as a regressor adjusts for the pre-trial level without modeling how the aperiodic background evolves within a trial. The same applies with more force to the pre-stimulus effect, which is elevated across the whole 3–30 Hz range rather than in any band. The difference spectrum of the pre-stimulus window (Figure 4F) is flat across the analyzed range, without the low-frequency weighting that shapes the post-stimulus difference, which is what a broadband shift in 1/f-like background power would produce. The spectral results should therefore be read as broadband power change of unresolved origin, not as evidence of theta or alpha modulation. Time-resolved parameterization would settle this, and the 7 s of clean pre-event data make the paradigm well suited to it.

### 4.3 Cardiac findings

Heart rate did not differentiate conditions at the group level in either window, and a classifier given cardiac features alone performed at chance in the same participants for whom the ERP was decodable above chance; adding those features to the EEG feature set changed nothing (Section 3.6, Supplementary Table S3). The null may reflect the low temporal resolution of RR-interval-derived heart rate relative to an event lasting one to two seconds, or a genuinely weak autonomic response to collisions in a passive VR context. Usable cardiac data in 14 of 18 participants nevertheless show that simultaneous cortical and cardiac recording in the headset is feasible.

### 4.4 Limitations

The study was ended at 18 of a planned 63 participants, so all results are exploratory, all effect sizes are provisional and likely inflated by the small sample, and no confirmatory inference is drawn. EDA and pupillometry, both preregistered as pre-stimulus measures, produced no usable data (Section 2.4); the pre-stimulus question is therefore addressed by EEG and cardiac data alone, and the modalities with the strongest prior support in this literature are absent.

The 12-channel montage with dry electrodes and no dedicated ocular reference limits spatial inference and constrains artifact correction, and component classification had to be handled by inspection rather than automatically. The oncoming vehicle also serves as the gaze target, so participants follow its approach from the far end of the scene to the blowout instead of fixating a cross. Pursuit and small saccades cannot be excluded from the pre-stimulus record, and without an eye tracker the time course of gaze is unobserved; the registered ocular and muscular control found no condition difference in peripheral power (Section 3.5), but a pursuit-related contribution to the pre-stimulus broadband effect remains possible at this sample size. Pupillometry, planned as the rigorous control for this limitation, was not available during data collection (Section 2.4).

The sample was older (mean 57 years), predominantly female, and reported high belief in intuition, which may limit generalization. Classification operated on participant-averaged responses with 32 observations, and 28 in the matched multimodal comparison; this is appropriate for the available sample and is tested across held-out participants, but the resulting confidence intervals are wide and the analysis cannot speak to single-trial decoding. The individual-difference analyses were uniformly null and at n = 14–16 cannot separate absence of moderation from insufficient sensitivity; they are reported only as hypothesis-generating.

## 5. Conclusion

In an immersive VR driving paradigm with quantum-randomized, temporally unpredictable collision events, evoked differentiation between collision and no-collision trials emerged after stimulus onset, converging across mass-univariate time-domain and multivariate decoding analyses. Before stimulus onset there was no time-domain effect and no decodable information, but a broadband time-frequency difference survived correction and passed every preregistered control analysis. We report it without interpreting it either way: the sample is small, the window was selected for being extreme, and one unreplicated cluster in one of three analysis domains is no basis for a positive claim; equally, it passed controls for the artifacts known to produce spurious pre-stimulus effects, so it cannot simply be dismissed. It is the clearest target this paradigm offers for adequately powered replication. Heart rate differentiated the conditions neither in the univariate model nor as classifier features, alone or combined with EEG. Because the study ended well short of its planned sample, these findings are exploratory: the post-stimulus effects require replication at their reported magnitude, and no pre-stimulus conclusion is drawn in either direction. What the study does establish is that a wireless, VR-integrated dry-electrode system can resolve robust event-related and oscillatory responses to naturalistic threat under conservative correction, and that a paradigm controlling the principal artifacts of pre-stimulus research is feasible and ready for adequately powered use.

## CRediT authorship contribution statement

Cédric Cannard: Conceptualization, Funding acquisition, Project administration, Resources, Methodology, Software, Investigation, Data curation, Formal analysis, Validation, Visualization, Writing – original draft, Writing – review & editing, Supervision. Demet Yeşilbaş: Methodology, Software (classification analyses), Formal analysis (classification analyses), Investigation, Data curation, Validation, Writing – review & editing.

## Ethics approval and consent

All study activities were approved and overseen by the Institutional Review Board (IRB) at the Institute of Noetic Sciences (IORG#0003743). The study was conducted in accordance with the principles embodied in the Declaration of Helsinki. All participants provided written informed consent prior to participation, including consent for their anonymized data to be shared openly, and received $30 compensation. The study was preregistered at https://osf.io/xuw34.

## Data availability statement

The analysis code, the per-participant classification analyses and the Brain Imaging Data Structure (BIDS) conversion pipeline are openly available at https://github.com/amisepa/galea-vr-driving-hazards (GPL-3.0); the EEGLAB plugin for importing and preprocessing recordings from this headset, with a step-by-step tutorial and sample data, is released separately at https://github.com/amisepa/galea-eeglab-plugin (version 1.7.1, GPL-3.0) and installable from the EEGLAB extension manager. The raw recordings are openly available in BIDS format with Hierarchical Event Descriptor annotations at OpenNeuro (https://openneuro.org/datasets/ds008837, version 1.1.3; doi: 10.18112/openneuro.ds008837.v1.1.3). The Unity VR application is available on request from the authors: it incorporates commercially licensed assets that do not permit redistribution, so it cannot be published openly. The delivered quantum-random-number trial sequences are included in the BIDS dataset events files.

## Declaration of generative AI and AI-assisted technologies in the manuscript preparation process

During the preparation of this work the authors used Claude (Anthropic, San Francisco, CA, USA) for language editing, formatting and software development assistance, and DeepSeek (DeepSeek, Hangzhou, China) for language editing and formatting. The authors directed all analyses and verified every reported result. After using these tools, the authors reviewed and edited the entire text and take full responsibility for the content of the published article.

## Supplementary Material

### S1. Quantum random number generator sequence diagnostics

Trial sequences were generated before data collection with a quantum random number generator and validated blind, before any recording took place. All 100 pre-generated sequences were tested; the table summarizes the four diagnostics across them. The full per-sequence values are in data/random_stim_sequences/qrandom_diagnostics_summary.csv in the repository.

**Table S1.** Randomization diagnostics across the 100 pre-generated trial sequences. Each sequence assigns 120 trials independently at 50% probability. Entropy is computed on the binary condition sequence, where 1.0 is the maximum for an equiprobable binary source. The runs test evaluates the null hypothesis that the sequence is independently ordered; autocorrelation is reported as the largest absolute coefficient over lags 1 to 10.

| **Diagnostic** | **Mean** | **SD** | **Minimum** | **Maximum** |
|---|---|---|---|---|
| Proportion of collision trials | 0.504 | 0.047 | 0.375 | 0.608 |
| Shannon entropy (bits) | 0.9937 | 0.0087 | 0.9544 | 1.0000 |
| Wald–Wolfowitz runs test, p | 0.514 | 0.278 | 0.003 | 1.000 |
| Maximum \|autocorrelation\|, lags 1–10 | 0.163 | 0.047 | 0.084 | 0.325 |

Of the 100 sequences, 5 returned a runs-test p below .05 (5%), which is what independence predicts at that threshold. No sequence was rejected or regenerated on the basis of these diagnostics; they are reported to document that the source behaved as a random binary generator before any data were collected.

### S2. Exploratory individual-difference models

Each individual-difference variable was entered separately as a second-level covariate in the hierarchical GLM, in the pre-stimulus and post-stimulus windows. Personality variables were available for 14 of the 16 participants and all other variables for all 16. Because each model was cluster-mass corrected within itself and none produced a surviving cluster, the registered false discovery rate step across variables had no rejections to adjust: Benjamini–Hochberg adjusted p-values are never smaller than the raw ones, so no family-level step could yield a rejection here.

These results are exploratory on grounds of statistical power and are not treated as findings of the study.

**Table S2a.** The 26 time-domain individual-difference models. Cluster-mass permutation correction, α = 0.05, 1000 permutations, as for the main analysis (Section 2.6). No model produced a cluster surviving correction in either window.

| Window | Covariate | n | Clusters surviving correction |
|---|---|---|---|
| Post-stimulus | age | 16 | 0 |
| Post-stimulus | sex | 16 | 0 |
| Post-stimulus | education | 16 | 0 |
| Post-stimulus | driving | 16 | 0 |
| Post-stimulus | sports | 16 | 0 |
| Post-stimulus | videogames | 16 | 0 |
| Post-stimulus | meditation | 16 | 0 |
| Post-stimulus | intuition | 16 | 0 |
| Post-stimulus | extraversion | 14 | 0 |
| Post-stimulus | agreeableness | 14 | 0 |
| Post-stimulus | conscientiousness | 14 | 0 |
| Post-stimulus | emotional stability | 14 | 0 |
| Post-stimulus | openness | 14 | 0 |
| Pre-stimulus | age | 16 | 0 |
| Pre-stimulus | sex | 16 | 0 |
| Pre-stimulus | education | 16 | 0 |
| Pre-stimulus | driving | 16 | 0 |
| Pre-stimulus | sports | 16 | 0 |
| Pre-stimulus | videogames | 16 | 0 |
| Pre-stimulus | meditation | 16 | 0 |
| Pre-stimulus | intuition | 16 | 0 |
| Pre-stimulus | extraversion | 14 | 0 |
| Pre-stimulus | agreeableness | 14 | 0 |
| Pre-stimulus | conscientiousness | 14 | 0 |
| Pre-stimulus | emotional stability | 14 | 0 |
| Pre-stimulus | openness | 14 | 0 |

Because the pre-stimulus effect exists only in the time-frequency domain, the time-domain models above cannot test the registered moderator hypothesis against it. Each participant’s condition effect was therefore summarized as mean power within each corrected time-frequency cluster, and each variable was correlated with it across participants in both windows. Benjamini– Hochberg was applied across all 26 tests, as registered. No association survives correction (smallest q = 0.81). The cluster was selected for its group-level condition effect, which inflates the cluster mean but does not bias a between-participant correlation, since the selection is blind to how individuals rank on any questionnaire.

**Table S2b.** The 26 time-frequency individual-difference correlations: Pearson r between each variable and the participant’s mean cluster power (collision minus no-collision, dB). q is the Benjamini–Hochberg adjusted p-value across all 26 tests.

| **Window** | **Covariate** | **n** | **r** | **p** | **q** |
|---|---|---|---|---|---|
| Pre-stimulus | age | 16 | -0.14 | 0.594 | 0.813 |
| Pre-stimulus | sex | 16 | -0.27 | 0.312 | 0.813 |
| Pre-stimulus | education | 16 | -0.22 | 0.405 | 0.813 |
| Pre-stimulus | driving | 16 | -0.17 | 0.535 | 0.813 |
| Pre-stimulus | sports | 16 | -0.25 | 0.343 | 0.813 |
| Pre-stimulus | videogames | 16 | 0.12 | 0.646 | 0.840 |
| Pre-stimulus | meditation | 16 | 0.06 | 0.832 | 0.901 |
| Pre-stimulus | intuition | 16 | -0.17 | 0.529 | 0.813 |
| Pre-stimulus | extraversion | 14 | -0.19 | 0.510 | 0.813 |
| Pre-stimulus | agreeableness | 14 | -0.21 | 0.477 | 0.813 |
| Pre-stimulus | conscientiousness | 14 | -0.16 | 0.581 | 0.813 |
| Pre-stimulus | emotional stability | 14 | 0.16 | 0.583 | 0.813 |
| Pre-stimulus | openness | 14 | 0.23 | 0.439 | 0.813 |
| Post-stimulus | age | 16 | -0.01 | 0.967 | 0.967 |
| Post-stimulus | sex | 16 | -0.40 | 0.129 | 0.813 |
| Post-stimulus | education | 16 | 0.07 | 0.800 | 0.901 |
| Post-stimulus | driving | 16 | -0.50 | 0.047 | 0.813 |
| Post-stimulus | sports | 16 | 0.09 | 0.735 | 0.901 |
| Post-stimulus | videogames | 16 | 0.08 | 0.774 | 0.901 |
| Post-stimulus | meditation | 16 | -0.29 | 0.279 | 0.813 |
| Post-stimulus | intuition | 16 | -0.38 | 0.143 | 0.813 |
| Post-stimulus | extraversion | 14 | 0.46 | 0.098 | 0.813 |
| Post-stimulus | agreeableness | 14 | -0.03 | 0.908 | 0.945 |
| Post-stimulus | conscientiousness | 14 | 0.36 | 0.211 | 0.813 |
| Post-stimulus | emotional stability | 14 | -0.23 | 0.424 | 0.813 |
| Post-stimulus | openness | 14 | 0.25 | 0.392 | 0.813 |

### S3. Electroencephalography, heart rate and their combination

The cardiac quality screen retained a different subset of participants from the EEG screen. To compare the modalities fairly, this analysis is restricted to the 14 participants with usable data in both, so that all three feature sets are evaluated on identical folds (28 observations). Cross-validation, classifiers and metrics are otherwise as in Section 2.8. The windows differ by modality: the event-related potential (ERP) features span the post-stimulus window (0 to +1200 ms), while the heart-rate features are mean instantaneous heart rate at eleven points spanning the full −5 to +5 s epoch. Heart rate yields roughly one sample per beat, so a 1.2 s window would hold only one or two values. The feature sets are therefore matched on participants and folds but not on time window.

**Table S3.** Classification of collision versus no-collision on the matched sample of 14 participants, using the post-stimulus ERP alone (0 to +1200 ms), eleven heart-rate values spanning the full −5 to +5 s epoch alone, and the two concatenated. Leave-one-subject-out cross-validation with predictions pooled across folds. All values are percentages except AUC. Chance accuracy is 50%.

| **Features** | **Classifier** | **Accuracy** | **95% CI** | **Sens.** | **Spec.** | **Prec.** | **F1** | **AUC** |
|---|---|---|---|---|---|---|---|---|
| EEG | SVM | 82.1 | [64, 92] | 78.6 | 85.7 | 84.6 | 81.5 | 0.85 |
| EEG | k-nearest neighbours | 78.6 | [60, 90] | 64.3 | 92.9 | 90.0 | 75.0 | 0.92 |
| EEG | Naive Bayes | 78.6 | [60, 90] | 78.6 | 78.6 | 78.6 | 78.6 | 0.85 |
| EEG | Random forest | 85.7 | [69, 94] | 78.6 | 92.9 | 91.7 | 84.6 | 0.91 |
| EEG | Decision tree | 82.1 | [64, 92] | 85.7 | 78.6 | 80.0 | 82.8 | 0.80 |
| EEG | Neural network (MLP) | 85.7 | [69, 94] | 71.4 | 100.0 | 100.0 | 83.3 | 0.90 |
| Heart rate | SVM | 50.0 | [33, 67] | 64.3 | 35.7 | 50.0 | 56.3 | 0.46 |
| Heart rate | k-nearest neighbours | 46.4 | [30, 64] | 21.4 | 71.4 | 42.9 | 28.6 | 0.48 |
| Heart rate | Naive Bayes | 50.0 | [33, 67] | 42.9 | 57.1 | 50.0 | 46.2 | 0.50 |
| Heart rate | Random forest | 46.4 | [30, 64] | 57.1 | 35.7 | 47.1 | 51.6 | 0.53 |
| Heart rate | Decision tree | 53.6 | [36, 70] | 35.7 | 71.4 | 55.6 | 43.5 | 0.53 |
| Heart rate | Neural network (MLP) | 50.0 | [33, 67] | 64.3 | 35.7 | 50.0 | 56.3 | 0.49 |
| EEG + heart rate | SVM | 82.1 | [64, 92] | 78.6 | 85.7 | 84.6 | 81.5 | 0.85 |
| EEG + heart rate | k-nearest neighbours | 78.6 | [60, 90] | 64.3 | 92.9 | 90.0 | 75.0 | 0.92 |
| EEG + heart rate | Naive Bayes | 78.6 | [60, 90] | 78.6 | 78.6 | 78.6 | 78.6 | 0.85 |
| EEG + heart rate | Random forest | 85.7 | [69, 94] | 85.7 | 85.7 | 85.7 | 85.7 | 0.96 |
| EEG + heart rate | Decision tree | 82.1 | [64, 92] | 85.7 | 78.6 | 80.0 | 82.8 | 0.80 |
| EEG + heart rate | Neural network (MLP) | 82.1 | [64, 92] | 71.4 | 92.9 | 90.9 | 80.0 | 0.90 |

Heart rate alone is at chance in every classifier, with all six confidence intervals (CIs) spanning 50% and areas under the curve (AUC) between 0.46 and 0.53. This matches the univariate cardiac analysis, which found no condition difference in event-related heart rate in either window. Adding the cardiac features to the ERP moves mean accuracy from 82.1% to 81.5%, and 4 of the 6 classifiers return predictions identical to the EEG-only model on every observation. That is what dimensionality alone predicts, and it is why the concatenation is not interpreted on its own: eleven features cannot displace several thousand. The heart-rate-only column is the interpretable one, and it indicates that the combination had nothing to gain.

### S4. Time-symmetry correlations on the time-frequency clusters

Registered hypothesis H4 held that the magnitude of pre-stimulus differentiation correlates positively with the magnitude of the post-stimulus response. The registered analysis is peak-matched to the primary mass-univariate effects; since the pre-stimulus effect was found in the time-frequency domain, trial-wise indices were taken as the mean channel-averaged power within the pre-stimulus and post-stimulus cluster windows (Section 2.7 of the main text), correlated per participant with the skipped Spearman estimator of the Robust Correlation Toolbox (Pernet et al., 2013), and tested against 10,000 within-participant permutations of the pre–post pairing. The group-level test is a one-sample t test on Fisher-transformed coefficients; the trimmed mean and its confidence interval are a bootstrap over participants (5,000 resamples). Because the windows come from these data, the analysis is exploratory.

**Table S4.** Per-participant time-symmetry correlations on the time-frequency clusters. r is the skipped Spearman correlation between trial-wise mean power in the pre-stimulus cluster window and in the post-stimulus cluster window; p is the two-tailed permutation p-value (10,000 pairing permutations). Trials are the retained artifact-free epochs entering each correlation.

| Participant | r | p (perm) | Trials | Collision | No-collision |
|---|---|---|---|---|---|
| sub-001 | 0.254 | 0.015 | 115 | 53 | 62 |
| sub-002 | 0.267 | 0.021 | 88 | 49 | 39 |
| sub-003 | 0.490 | < .001 | 106 | 46 | 60 |
| sub-004 | 0.242 | 0.034 | 104 | 49 | 55 |
| sub-005 | 0.207 | 0.046 | 115 | 56 | 59 |
| sub-006 | 0.255 | 0.016 | 114 | 50 | 64 |
| sub-007 | 0.238 | 0.029 | 113 | 56 | 57 |
| sub-008 | 0.467 | < .001 | 106 | 55 | 51 |
| sub-009 | 0.231 | 0.053 | 88 | 53 | 35 |
| sub-010 | 0.257 | 0.032 | 94 | 45 | 49 |
| sub-011 | 0.184 | 0.099 | 108 | 50 | 58 |
| sub-012 | 0.282 | 0.019 | 91 | 42 | 49 |
| sub-013 | 0.799 | < .001 | 94 | 55 | 39 |
| sub-014 | 0.264 | 0.017 | 106 | 53 | 53 |
| sub-015 | 0.278 | 0.027 | 86 | 49 | 37 |
| sub-016 | 0.435 | < .001 | 107 | 43 | 64 |

**Figure S1.**
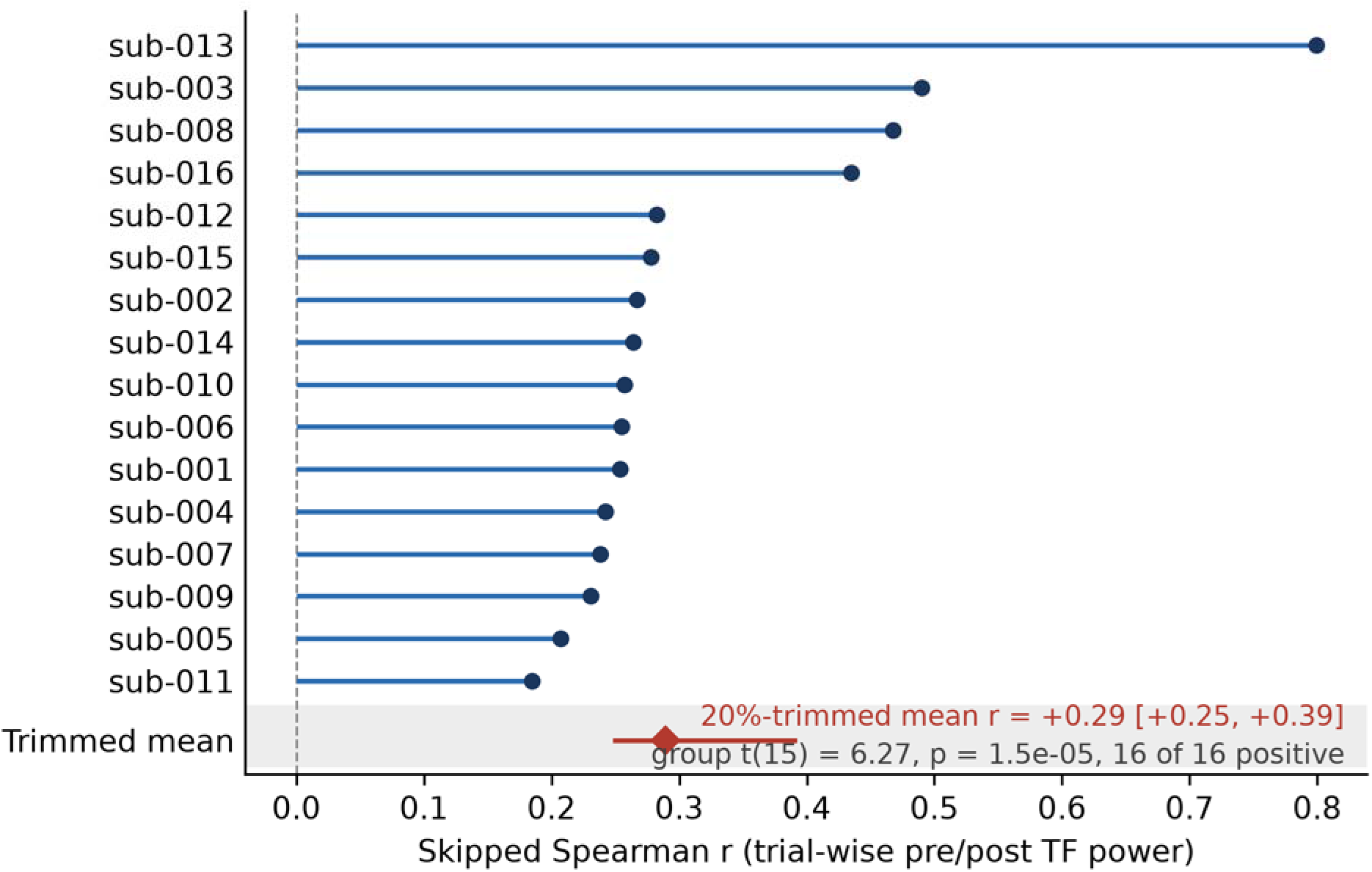
Per-participant time-symmetry correlations (skipped Spearman r between trial-wise pre- and post-stimulus power in the two time-frequency cluster windows), sorted by magnitude. The red diamond marks the group-level 20%-trimmed mean with its bootstrap 95% confidence interval; all 16 of 16 participants are positive.

